## Supplementary material for "Evolutionary implications of the first microRNA- and piRNA complement of *Lepidodermella squamata* (Gastrotricha)": ST1

Supplementary table S1: Curated set of 65 RNA proteins from *C. elegans* used for reciprocal Blast searches.

| Gene | *C. elegans* gene ID  WS264 release | RNAi complex | Protein name |
| --- | --- | --- | --- |
| *AIN-1* | C06G1.4 | RNA Induced Silencing | Alg-1 INteracting protein-1/GW182 |
| *AIN-2* | B0041.2a | RNA Induced Silencing | Alg-2 INteracting protein-2/GW182 |
| *ALG-1* | F48F7.1b | RNA Induced Silencing | Argonaute like gene-1 |
| *ALG-2* | T07D3.7a | RNA Induced Silencing | Argonaute like gene-2 |
| *CGH-1* | C07H6.5 | RNA Induced Silencing | Conserved germline helicase |
| *GLD-1* | T23G11.3 | RNA Induced Silencing | Defective in germ line development |
| *ASD-2* | T21G5.5a | RNA Induced Silencing | Alternative splicing defective |
| *NHL-2* | F26F4.7 | RNA Induced Silencing | NHL domain containing |
| *TSN-1* | F17G7.2 | RNA Induced Silencing | Tudor similar nuclease |
| *VIG-1* | F56D12.5a | RNA Induced Silencing | Vasa intronic gene-1 |
| *RDE-1* | K08H10.7 | RNA Induced Silencing | RNAi deficient-1 |
| *XRN-1* | Y39G8C.1 | RNA Induced Silencing | XRN ribonuclease related |
| *SID-1* | C04F5.1 | Systemic RNA Interference | Systemic RNA interference defective-1 |
| *SID-2* | ZK520.2 | Systemic RNA Interference | Systemic RNA interference defective-2 |
| *RSD-2* | F52G2.2a | Systemic RNA Interference | RNA spreading defective-2 |
| *RSD-3* | C34E11.1 | Systemic RNA Interference | RNA spreading defective-3 |
| *RSD-6* | F16D3.2 | Systemic RNA Interference | RNA spreading defective-6 |
| *DCR-1* | K12H4.8 | Dicer | Dicer-1 |
| *DRH-1* | F15B10.2 | Dicer | Dicer related helicase-1 |
| *RDE-4* | T20G5.11 | Dicer | RNAi decient-4 |
| *DRSH-1* | F26E4.10a | Microprocessor | Drosha-1 |
| *PASH-1* | T22A3.5a | Microprocessor | Pasha-1 |
| *DRH-3* | D2005.5 | ERI | Dicer related helicase-3 |
| *ERGO-1* | R09A1.1a | ERI | Endogenous RNAi deficient arGOnaute |
| *ERI-1* | T07A9.5a | ERI | Enhanced RnaI-1 |
| *ERI-3* | W09B6.3a | ERI | Enhanced RnaI-3 |
| *ERI-5* | Y38F2AR.1a | ERI | Enhanced RnaI-5 |
| *ERI-6* | C41D11.1a | ERI | Enhanced RnaI-6 |
| *ERI-7* | C41D11.7 | ERI | Enhanced RnaI-7 |
| *RRF-3* | F10B5.7 | ERI | RNA dependent polymerase family |
| *R02D3.8* | R02D3.8 | ERI | Hypothetical protein (paralog to ERI-1) |
| *WAGO-4* | F58G1.1 | Secondary Argonautes | Worm ArGOnaute 4 |
| *PPW-1* | C18E3.7c | Secondary Argonautes | PAZ/PIWI domain containing-1 |
| *PPW-2* | Y110A7A.18 | Secondary Argonautes | PAZ/PIWI domain containing-2 |
| *SAGO-1* | K12B6.1 | Secondary Argonautes | Synthetic secondary siRNA-deficient ArGOnaute mutant-1 |
| *SAGO-2* | F56A6.1a | Secondary Argonautes | Synthetic secondary siRNA-deficient ArGOnaute mutant-2 |
| *EGO-1* | F26A3.3 | RNA Dependent RNA Polymerases | Enhancer of Glp-One |
| *PIR-1* | T23G7.5a | RNA Dependent RNA Polymerases | Phosphatase interacting with RNA/RNP |
| *RRF-1* | F26A3.8a | RNA Dependent RNA Polymerases | RNA dependent polymerase family |
| *RRF-2* | M01G12.12 | RNA Dependent RNA Polymerases | RNA dependent polymerase family |
| *CSR-1* | F20D12.1a | Secondary RNAi Silencing | Chromosome segregation and RNAi deficient |
| *MUT-14* | C14C11.6 | Secondary RNAi Silencing | MUTator-14 |
| *MUT-16* | B0379.3a | Secondary RNAi Silencing | MUTator-16 |
| *RDE-3* | K04F10.6a | Secondary RNAi Silencing | RNAi deficient-3 |
| *MES-2* | R06A4.7 | Polycomb | Maternal effect sterile-2 |
| *MES-3* | F54C1.3a | Polycomb | Material effect sterile-3 |
| *MES-6* | C09G4.5 | Polycomb | Material effect sterile-6 |
| *XPO-1* | ZK742.1a | Others | eXPOrtin |
| *MUT-7* | ZK1098.8 | Others | MUTator-7 |
| *RDE-2* | F21C3.4a | Others | RNAi deficient-2 |
| *GLD-2* | ZC308.1a | Others | Poly(A) RNA polymerase (homolog to RDE-3) |
| *SMG-2* | Y48G8AL.6 | Others | Suppresor with morphological effect on genitalia-2 |
| *SMG-5* | W02D3.8 | Others | Suppresor with morphological effect on genitalia-5 |
| *SMG-6* | Y54F10AL.2a | Others | Suppresor with morphological effect on genitalia-6 |
| *GFL-1* | M04B2.3 | Others | Human GAS41 like |
| *HENN-1* | C02F5.6b | Others | HEsitatioN behaviour |
| *HPL-1* | K08H2.6 | Others | Heterochromatin protein like-1 |
| *MRG-1* | Y37D8A.9a | Others | Mortality factor related gene |
| *NRDE-3* | R04A9.2 | Others | Nuclear RNAi defective-3 |
| *PRMT-5* | C34E10.5 | Others | Protein arginine Methil Transferase |
| *RHA-1* | T04D4.3 | Others | Rna HelicAse |
| *PHF-14* | Y59A8A.2 | Others | PHd finger-14 (paralog to ZFP-1) |
| *PHF-15* | Y53G8AR.2a | Others | PHd finger-15 (paralog to ZFP-1) |
| *LIN-49* | F42A9.2 | Others | Bromodomain containing protein (paralog to ZFP-1) |
| *ZFP-1* | F54F2.2a | Others | Zinc finger protein |
