## Supplementary material for "Evolutionary implications of the first microRNA- and piRNA complement of *Lepidodermella squamata* (Gastrotricha)": ST3

Supplementary table S3: PIWI-like proteins identified in metazoan predicted proteins and gastrotrich transcriptomes

| Brachiopoda | |
| --- | --- |
| *Lingula anatina* | Lana.g23732.t1, Lana.g1407.t1, Lana.g15951.t1 |
| **Phoronida** | |
| *Phoronis australis* | Paus.g1500.t1 |
| **Annelida** | |
| *Capitella teleta* | Ctel.CapteP163584, Ctel.CapteP154759 |
| *Helobdella robusta* | Hrob.HelroP75625, Hrob.HelroP65566 |
| **Nemertea** | |
| *Notospermus geniculatus* | Ngen.g6114.t1, Ngen.g12035.t1, Ngen.g13861.t1, Ngen.g39228.t1, Ngen.g40598.t1 |
| **Mollusca** | |
| *Aplysia californica* | Acal.XP_005096149.1, Acal.XP_012940110.1 |
| *Argopecten purpuratus* | Apur.evm.model.scaffold_633.50_evm.model.scaffold_633.51, Apur.evm.model.scaffold_53598.11 |
| *Bathymodiolus platifrons* | Bpla.Bpl_scaf_60737-1.14, Bpla.Bpl_scaf_45124-0.37, Bpla.Bpl_scaf_60737-0.5 |
| *Biomphalaria glabrata* | Bgla.BGLB010170-PA |
| *Chlamys farreri* | Cfar.CF10513.1, Cfar.CF64453.3 |
| *Crassostrea gigas* | Cgig.EKC35279, Cgig.EKC29295 |
| *Haliotis discus hannai* | Hdih.HDSC07612CG00020 |
| *Lottia gigantea* | Lgig.LotgiP131825, Lgig.LotgiP210915 |
| *Modiolus philippinarum* |  |
| *Patinopecten yessoensis* | Pyes.PY_T05166, Pyes.PY_T16374 |
| *Pinctada fucata* | Pfuc.pfu_aug2.0_267.1_23780.t1 |
| *Pinctada fucata martensii* | Pfum.Pma_10020092 |
| **Gastrotricha** | |
| *Dactylopodola baltica* |  |
| *Diuronotus aspetos* | Dasp.Gene.108664 |
| *Lepidodermella squamata* | Lsqu.Gene.78021, Lsqu.Gene.176266, Lsqu.Gene.20027, Lsqu.Gene.203941, Lsqu.Gene.95003 |
| *Macrodasys sp.* |  |
| *Megadasys sp.* | Mega.Gene.23503 |
| *Mesodasys laticaudatus* |  |
| **Platyhelminthes** | |
| *Echinococcus multilocularis* | Emul.EmuJ_000739100.1 |
| *Hymenolepis microstoma* | Hmic.HmN_002103300.1 |
| *Macrostomum lignano* | Mlig.maker-uti_cns_0001573-snap-gene-0.4-mRNA-1, Mlig.maker-uti_cns_0001575-snap-gene-0.7-mRNA-1, Mlig.maker-uti_cns_0003023-snap-gene-0.7-mRNA-1, Mlig.maker-uti_cns_0003083-snap-gene-0.18-mRNA-1, Mlig.maker-uti_cns_0009213-snap-gene-0.4-mRNA-1, Mlig.maker-uti_cns_0011474-snap-gene-0.6-mRNA-1, Mlig.maker-uti_cns_0047331-snap-gene-0.2-mRNA-1, Mlig.maker-uti_cns_0045470-snap-gene-0.10-mRNA-1, Mlig.maker-uti_cns_0045747-snap-gene-1.12-mRNA-1, Mlig.maker-uti_cns_0047984-snap-gene-0.6-mRNA-1 |
| *Schistosoma mansoni* |  |
| *Schmidtea mediterranea* | Smed.mk4.000945.06, Smed.mk4.000381.00, Smed.mk4.003473.02, Smed.mk4.001991.01, Smed.mk4.003182.00, Smed.mk4.002396.00, Smed.mk4.000678.05, Smed.mk4.001253.02, Smed.mk4.010770.02 |
| **Orthonectida** | |
| *Intoshia linei* | Ilin.OAF65019.1 |
| **Rotifera** | |
| *Adineta vaga* | Avag.GSADVT00033948001, Avag.GSADVT00030821001, Avag.GSADVT00039413001, Avag.GSADVT00010852001, Avag.GSADVT00010869001, Avag.GSADVT00065605001, Avag.GSADVT00052987001, Avag.GSADVT00036072001, Avag.GSADVT00043793001 |
| **Arthropoda** | |
| *Acyrthosiphon pisum* | Apis.ACYPI37002-PA, Apis.ACYPI005423-PA, Apis.ACYPI008719-PA, Apis.ACYPI008078-PA, Apis.ACYPI004068-PA, Apis.ACYPI005740-PA, Apis.ACYPI004735-PA, Apis.ACYPI008672-PA, Apis.ACYPI062374-PA |
| *Anopheles gambiae* | Agam.AGAP011204-PA, Agam.AGAP008862-PA |
| *Apis mellifera* | Amel.gnl\|Amel_4.5\|GB49909-PA |
| *Culex quinquefasciatus* | Cqui.CPIJ012516-PA, Cqui.CPIJ005275-PA, Cqui.CPIJ017381-PA |
| *Daphnia pulex* | Dpul.EFX83176, Dpul.EFX83177, Dpul.EFX83174, Dpul.EFX83175, Dpul.EFX88764, Dpul.EFX71843, Dpul.EFX71842, Dpul.EFX81763 |
| *Drosophila melanogaster* | Dmel.FBpp0309202, Dmel.FBpp0079755, Dmel.FBpp0290939, Dmel.FBpp0289159, Dmel.FBpp0289158, Dmel.FBpp0289160 |
| *Heliconius melpomene* | Hmel.HMEL007215-PA, Hmel.HMEL008054-PA |
| *Ixodes scapularis* |  |
| *Nasonia vitripennis* | Nvit.NV10813-PA, Nvit.NV12150-PA, Nvit.NV12181-PA |
| *Pediculus humanus* | Phum.PHUM411830-PA |
| *Strigamia maritima* |  |
| *Tribolium castaneum* |  |
| **Tardigrada** | |
| *Hysibius dujardini* | Hduj.nHd.2.3.1.t02500-RA, Hduj.nHd.2.3.1.t05118-RA, Hduj.nHd.2.3.1.t05707-RA, Hduj.nHd.2.3.1.t08161-RA, Hduj.nHd.2.3.1.t10523-RA, Hduj.nHd.2.3.1.t13281-RA |
| *Ramazzottius varieomatus* | Rvar.RVARI.g9391.t1, Rvar.RVARI.g10304.t1, Rvar.RVARI.g10741.t1, Rvar.RVARI.g11451.t1 |
| **Nematoda** | |
| *Brugia malayi* | Bmal.Bm4557, Bmal.Bm5200a, Bmal.Bm5200b |
| *Caenorhabditis elegans* | Cele.C18E3.7d, Cele.D2030.6, Cele.C01G5.2a, Cele.C14B1.7a, Cele.C16C10.3, Cele.F58G1.1, Cele.F55A12.1, Cele.K12B6.1, Cele.R04A9.2, Cele.R09A1.1a, Cele.R09A1.1b, Cele.T22H9.3, Cele.Y49F6A.1, Cele.ZK1248.7 |
| *Loa loa* | Lloa.EJD75956.1, Lloa.EFO28277.2 |
| *Onchocerca volvulus* | Ovol.OVOC1650, Ovol.OVOC10749 |
| *Pristionchus pacificus* | Ppac.PPA07002, Ppac.PPA09273, Ppac.PPA09363, Ppac.PPA11660, Ppac.PPA17322, Ppac.PPA22662, Ppac.PPA23639, Ppac.PPA29011 |
| *Strongyloides ratti* | Srat.SRAE_1000016100, Srat.SRAE_1000023200, Srat.SRAE_1000116200, Srat.SRAE_1000217900 |
| *Trichinella spiralis* |  |
| **Deuterostomia** | |
| *Acanthaster planci* | Apla.gbr.2.28.t1, Apla.gbr.260.7.t1 |
| *Anolis carolinensis* |  |
| *Branchiostoma floridae* | Bflo.jgi\|Brafl1\|253549\|e_gw.649.3.1, Bflo.jgi\|Brafl1\|89999\|fgenesh2_pg.scaffold_193000004, Bflo.jgi\|Brafl1\|91808\|fgenesh2_pg.scaffold_214000001, Bflo.jgi\|Brafl1\|126411\|estExt_fgenesh2_pg.C_2140014 |
| *Ciona intestinalis* | Cint.jgi\|Cioin2\|261286\|gw1.475.2.1 |
| *Danio rerio* |  |
| *Gallus gallus* |  |
| *Homo sapiens* | Hsap.ENSP00000431843.1, Hsap.ENSP00000435718.1, Hsap.ENSP00000330031.5, Hsap.ENSP00000479524.1 |
| *Loxodonta africana* | Lafr.ENSLAFP00000007066.4 |
| *Monodelphis domestica* | Mdom.ENSMODP00000000258.2 |
| *Oikopleura dioica* | Odio.GSOIDP00004070001, Odio.GSOIDP00012941001 |
| *Ornithorhynchus anatinus* |  |
| *Petromyzon marinus* | Pmar.ENSPMAP00000001860.1 |
| *Ptychodera flava* | Pfla.PFL3_pfl_40v0_9_20150316_1g2512.t1, Pfla.PFL3_pfl_40v0_9_20150316_1g27269.t1 |
| *Saccoglossus kowalevskii* | Skow.Sakowv30028355m, Skow.Sakowv30008752m |
| *Strongylocentrotus purpuratus* |  |
| *Tursiops truncatus* |  |
| *Xenopus tropicalis* | Xtro.ENSXETP00000027524.3, Xtro.ENSXETP00000014018.2 |
| **Non-Bilateria** | |
| *Acropora digitifera* | Adig.adi_v1.20994, Adig.adi_v1.13833 |
| *Amphimedon queenslandica* | Aque.PAC:15726897 |
| *Hydra magnipapillata* |  |
| *Leucosolenia complicata* | Lcom.lcpid33121\|, Lcom.lcpid86122\| |
| *Mnemiopsis leidyi* |  |
| *Nematostella vectensis* | Nvec.EDO49931, Nvec.EDO34027 |
| *Oscarella carmela* | Ocar.m.25027 |
| *Pleurobranchia bachei* | Pbac.sb\|3462598\| |
| *Sycon ciliatum* | Scil.scpid38180\| |
| *Trichoplax adhaerens* |  |
