## Supplementary material for "Evolutionary implications of the first microRNA- and piRNA complement of *Lepidodermella squamata* (Gastrotricha)": ST4

Supplementary table S4: List of metazoan genomes used for RNAi protein survey.

| Species | Tag code | Phylogenetic group | Source |
| --- | --- | --- | --- |
| *Lingula anatina* | Lana | Brachipoda | Ensembl Metazoa Release 38 |
| *Phoronis australis* | Paus | Phoronida | OIST Marine Genomics Unit |
| *Capitelle teleta* | Ctel | Annelida | Ensembl Metazoa Release 38 |
| *Helobdella robusta* | Hrob | Annelida | Ensembl Metazoa Release 38 |
| *Notospermus geniculatus* | Ngen | Nemertea | OIST Marine Genomics Unit |
| *Aplysia californica* | Acal | Mollusca | NCBI Genome Assembly |
| *Argopecten purpuratus* | Apur | Mollusca | GigaScience |
| *Bathymodiolus platifrons* | Bpla | Mollusca | DataDryad |
| *Biomplaharia glabrata* | Bgla | Mollusca | VectorBase |
| *Chlamys farreri* | Cfar | Mollusca | CfBase |
| *Crassostrea gigas* | Cgig | Mollusca | Ensembl Metazoa Release 38 |
| *Haliotis discus hannai* | Hdih | Mollusca | GigaScience |
| *Lottia gigantea* | Lgig | Mollusca | Ensembl Metazoa Release 38 |
| *Modiolus philippinarum* | Mphi | Mollusca | DataDryad |
| *Patinopecten yessoensis* | Pyes | Mollusca | PyBase |
| *Pinctada fucata* | Pfuc | Mollusca | OIST Marine Genomics Unit |
| *Pinctada fucata martensii* | Pfum | Mollusca | GigaScience |
| *Echinococcus multilocularis* | Emul | Platyhelminthes | WormBase ParaSite WBPS9 |
| *Hymenolepis microstoma* | Hmic | Platyhelminthes | WormBase ParaSite WBPS9 |
| *Macrostomum lignano* | Mlig | Platyhelminthes | WormBase ParaSite WBPS9 |
| *Schistosoma mansoni* | Sman | Platyhelminthes | WormBase ParaSite WBPS9 |
| *Schmidtea mediterranea* | Smed | Platyhelminthes | WormBase ParaSite WBPS9 |
| *Intoshia linei* | Ilin | Orthonectida | NCBI Genome Assembly |
| *Adineta vaga* | Avag | Rotifera | Ensembl Metazoa Release 38 |
| *Acyrthosiphon pisum* | Apis | Arthropoda | Ensembl Metazoa Release 38 |
| *Anopheles gambiae* | Agam | Arthropoda | Ensembl Metazoa Release 38 |
| *Apis mellifera* | Amel | Arthropoda | BeeBase |
| *Culex quinquefasciatus* | Cqui | Arthropoda | Ensembl Metazoa Release 38 |
| *Daphnia pulex* | Dpul | Arthropoda | Ensembl Metazoa Release 38 |
| *Drosophila melanogaster* | Dmel | Arthropoda | Ensembl Metazoa Release 38 |
| *Heliconius melpomene* | Hmel | Arthropoda | Ensembl Metazoa Release 38 |
| *Ixodes scapularis* | Isca | Arthropoda | Ensembl Metazoa Release 38 |
| *Nasonia vitripennis* | Nvit | Arthropoda | Ensembl Metazoa Release 38 |
| *Pediculus humanus* | Phum | Arthropoda | Ensembl Metazoa Release 38 |
| *Strigamia maritima* | Smar | Arthropoda | Ensembl Metazoa Release 38 |
| *Tribolium castaneum* | Tcas | Arthropoda | Ensembl Metazoa Release 38 |
| *Hysibius dujardini* | Hduj | Tardigrada | tardigrades.org |
| *Ramazzottius varieomatus* | Rvar | Tardigrada | tardigrades.org |
| *Brugia malayi* | Bmal | Nematoda | Ensembl Metazoa Release 38 |
| *Caenorhabditis elegans* | Cele | Nematoda | Ensembl Metazoa Release 38 |
| *Loa loa* | Lloa | Nematoda | Ensembl Metazoa Release 38 |
| *Onchocerca volvulus* | Ovol | Nematoda | Ensembl Metazoa Release 38 |
| *Pristionchus pacificus* | Ppac | Nematoda | Ensembl Metazoa Release 38 |
| *Strongyloides ratti* | Srat | Nematoda | Ensembl Metazoa Release 38 |
| *Trichinella spiralis* | Tspi | Nematoda | Ensembl Metazoa Release 38 |
| *Acanthaster planci* | Apla | Deuterostomia | OIST Marine Genomics Unit |
| *Anolis carolinensis* | Acar | Deuterostomia | Ensembl Vertebrate Release 91 |
| *Branchiostoma floridae* | Bflo | Deuterostomia | Joint Genome Institute |
| *Ciona intestinalis* | Cint | Deuterostomia | Joint Genome Insitute |
| *Danio rerio* | Drer | Deuterostomia | Ensembl Vertebrate Release 91 |
| *Gallus gallus* | Ggal | Deuterostomia | Ensembl Vertebrate Release 91 |
| *Homo sapiens* | Hsap | Deuterostomia | Ensembl Vertebrate Release 91 |
| *Loxodonta africana* | Lafr | Deuterostomia | Ensembl Vertebrate Release 91 |
| *Monodelphis domestica* | Mdom | Deuterostomia | Ensembl Vertebrate Release 91 |
| *Oikopleura dioica* | Odio | Deuterostomia | OikoBase |
| *Ornithorhynchus anatinus* | Oana | Deuterostomia | Ensembl Vertebrate Release 91 |
| *Petromyzon marinus* | Pmar | Deuterostomia | Ensembl Vertebrate Release 91 |
| *Ptychodera flava* | Pfla | Deuterostomia | OIST Marine Genomics Unit |
| *Saccoglossus kowalevskii* | Skow | Deuterostomia | OIST Marine Genomics Unit |
| *Strongylocentrotus purpuratus* | Spur | Deuterostomia | Ensembl Metazoa Release 38 |
| *Tursiops truncates* | Stru | Deuterostomia | Ensembl Vertebrate Release 91 |
| *Xenopus tropicalis* | Xtro | Deuterostomia | Ensembl Vertebrate Release 91 |
| *Acropora digitifera* | Adig | Non-Bilateria | Compagen |
| *Amphimedon queenslandica* | Aque | Non-Bilateria | Ensembl Metazoa Release 38 |
| *Hydra magnipapillata* | Hmag | Non-Bilateria | Compagen |
| *Leucosolenia complicate* | Lcom | Non-Bilateria | Compagen |
| *Mnemiopsis leidyi* | Mlei | Non-Bilateria | Ensembl Metazoa Release 38 |
| *Nematostella vectensis* | Nvec | Non-Bilateria | Ensembl Metazoa Release 38 |
| *Oscarella carmela* | Ocar | Non-Bilateria | Compagen |
| *Pleurobranchia bachei* | Pbac | Non-Bilateria | Neurobase |
| *Sycon ciliatum* | Scil | Non-Bilateria | Compagen |
| *Trichoplax adhaerens* | Tadh | Non-Bilateria | Ensembl Metazoa Release 38 |
