## Supplementary material for "Evolutionary implications of the first microRNA- and piRNA complement of *Lepidodermella squamata* (Gastrotricha)": ST5

Supplementary table S5: List of gastrotrich transcriptomes used for RNAi protein survey.

| Species | Tag code | Phylogenetic group | NCBI SRA accession number |
| --- | --- | --- | --- |
| *Dactylopodola baltica* | Dbal | Gastrotricha | SRR1275388, SRR1275389, SRR1273672, SRR1273673 |
| *Diuronotus aspetos* | Dasp | Gastrotricha | SRR2131262 |
| *Lepidodermella squamata* | Lsqu | Gastrotricha | SRR1982110 |
| *Macrodasys sp.* | Macr | Gastrotricha | SRR1271706, SRR1271707, SRR1271708, SRR1275393 |
| *Megadasys sp.* | Mega | Gastrotricha | SRR1273711, SRR1273712, SRR1275394, SRR1275397 |
| *Mesodays laticaudatus* | Mlat | Gastrotricha | SRR1797883 |
