## Supplementary material for "Evolutionary implications of the first microRNA- and piRNA complement of *Lepidodermella squamata* (Gastrotricha)": ST6

Supplementary table S6: List of protostome genomes and gastrotrich transcriptomes used for PIWI-like phylogenetic analysis.

| Species | Phylogenetic group | Genome/Transcriptome |
| --- | --- | --- |
| *Lingula anatina* | Brachiopoda | Genome |
| *Phoronis australis* | Phoronida | Genome |
| *Capitella teleta* | Annelida | Genome |
| *Helobdella robusta* | Annelida | Genome |
| *Notospermus geniculatus* | Nemertea | Genome |
| *Aplysia californica* | Mollusca | Genome |
| *Argopecten purpuratus* | Mollusca | Genome |
| *Bathymodiolus platifrons* | Mollusca | Genome |
| *Biomplaharia glabrata* | Mollusca | Genome |
| *Chlamys farreri* | Mollusca | Genome |
| *Crassostrea gigas* | Mollusca | Genome |
| *Haliotis discus hannai* | Mollusca | Genome |
| *Lottia gigantea* | Mollusca | Genome |
| *Patinopecten yessoensis* | Mollusca | Genome |
| *Pinctada fucata* | Mollusca | Genome |
| *Pinctada fucata martensii* | Mollusca | Genome |
| *Diuronotus aspetos* | Gastrotricha | Transcriptome |
| *Lepidodermella squamata* | Gastrotricha | Transcriptome |
| *Megadasys sp.* | Gastrotricha | Transcriptome |
| *Echinococcus multilocularis* | Platyhelminthes | Genome |
| *Hymenolepis microstoma* | Platyhelminthes | Genome |
| *Macrostomum lignano* | Platyhelminthes | Genome |
| *Schmidtea mediterranea* | Platyhelminthes | Genome |
| *Intoshia linei* | Orthonectida | Genome |
