## Supplementary figures and images for "Evolutionary implications of the first microRNA- and piRNA complement of *Lepidodermella squamata* (Gastrotricha)"

### SF1

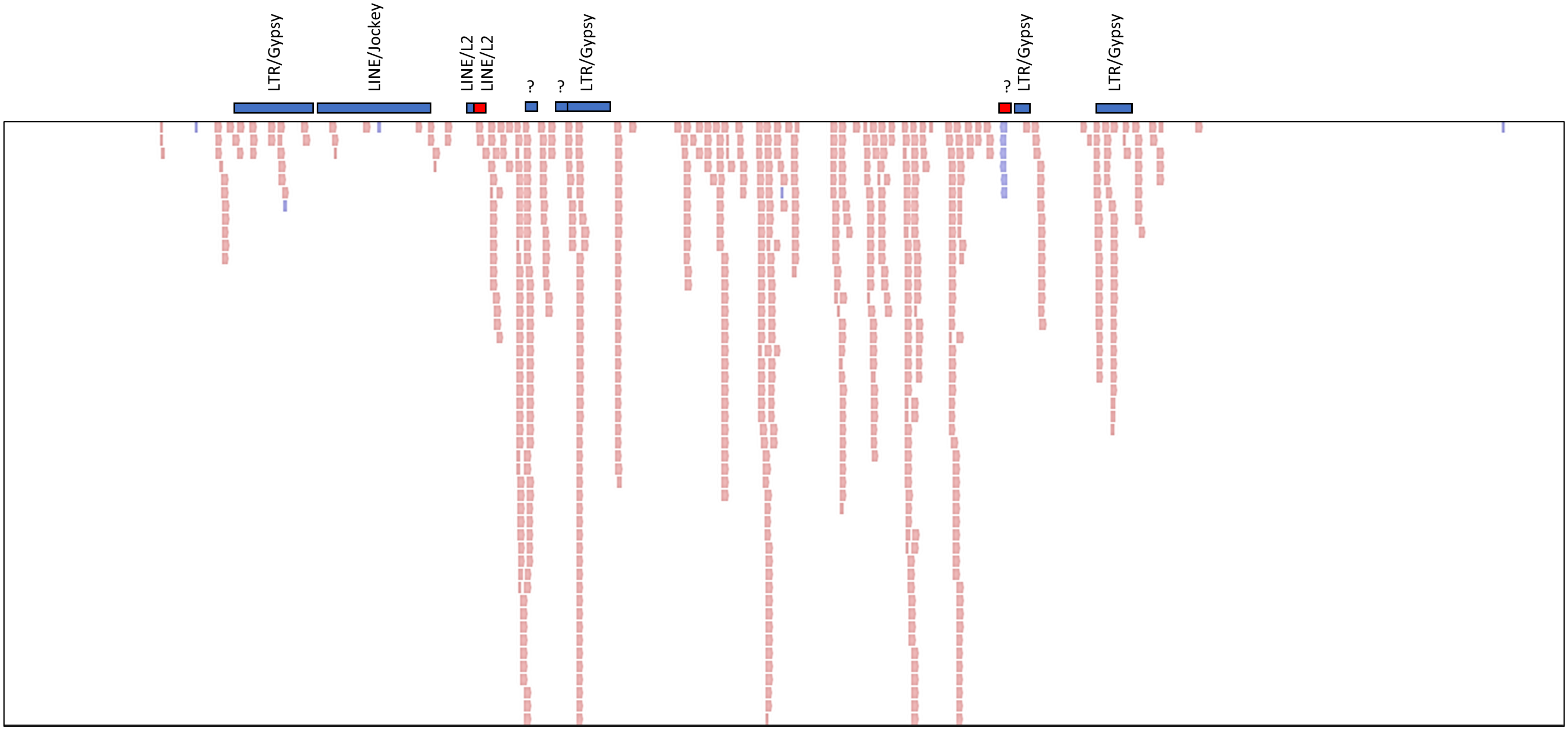


piRNA cluster #1: scaffold95_size236010:16,013-20,557

### SF2

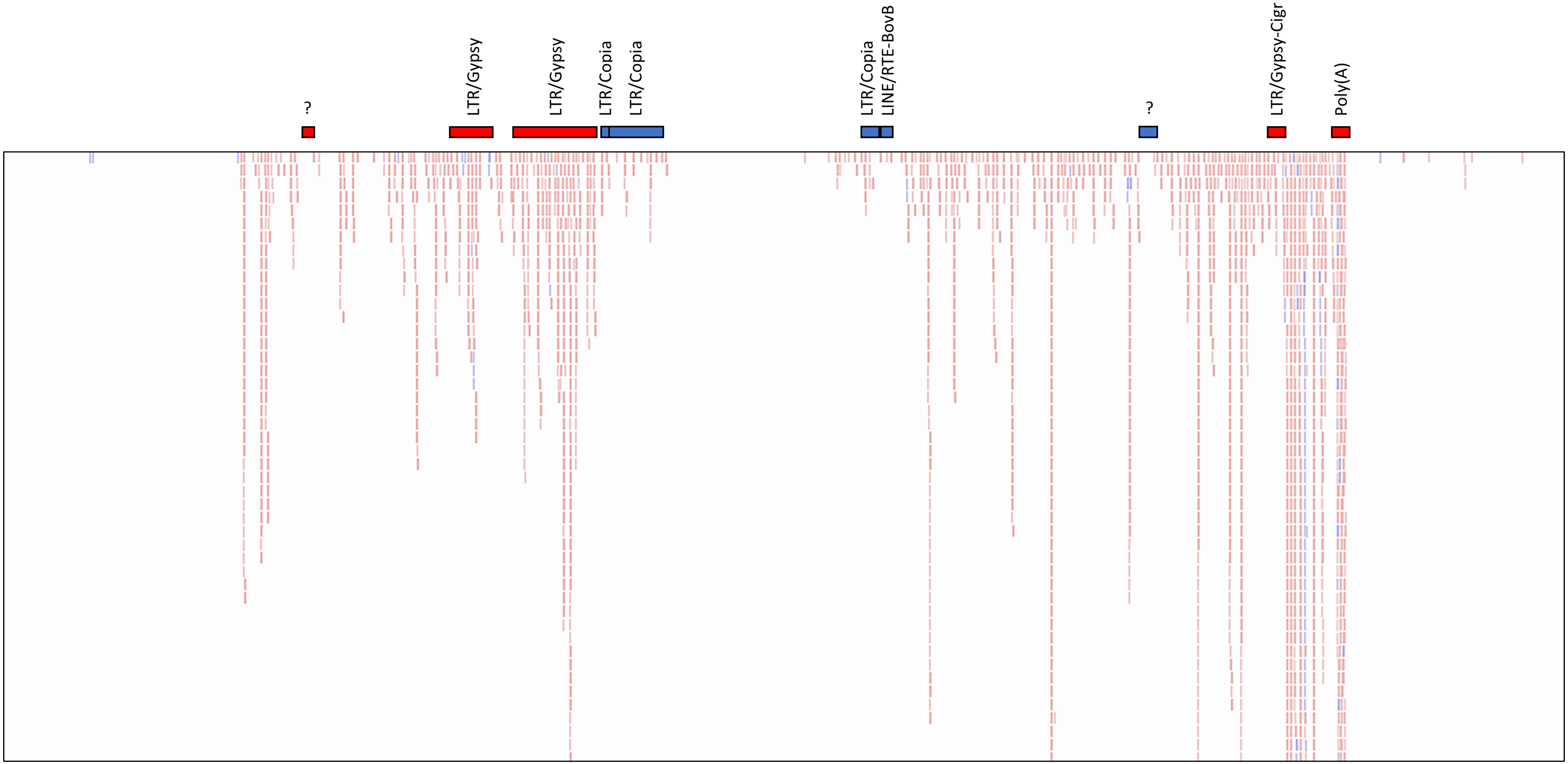


piRNA cluster #2: scaffold171_size172051:144,160-158,000

### SF3

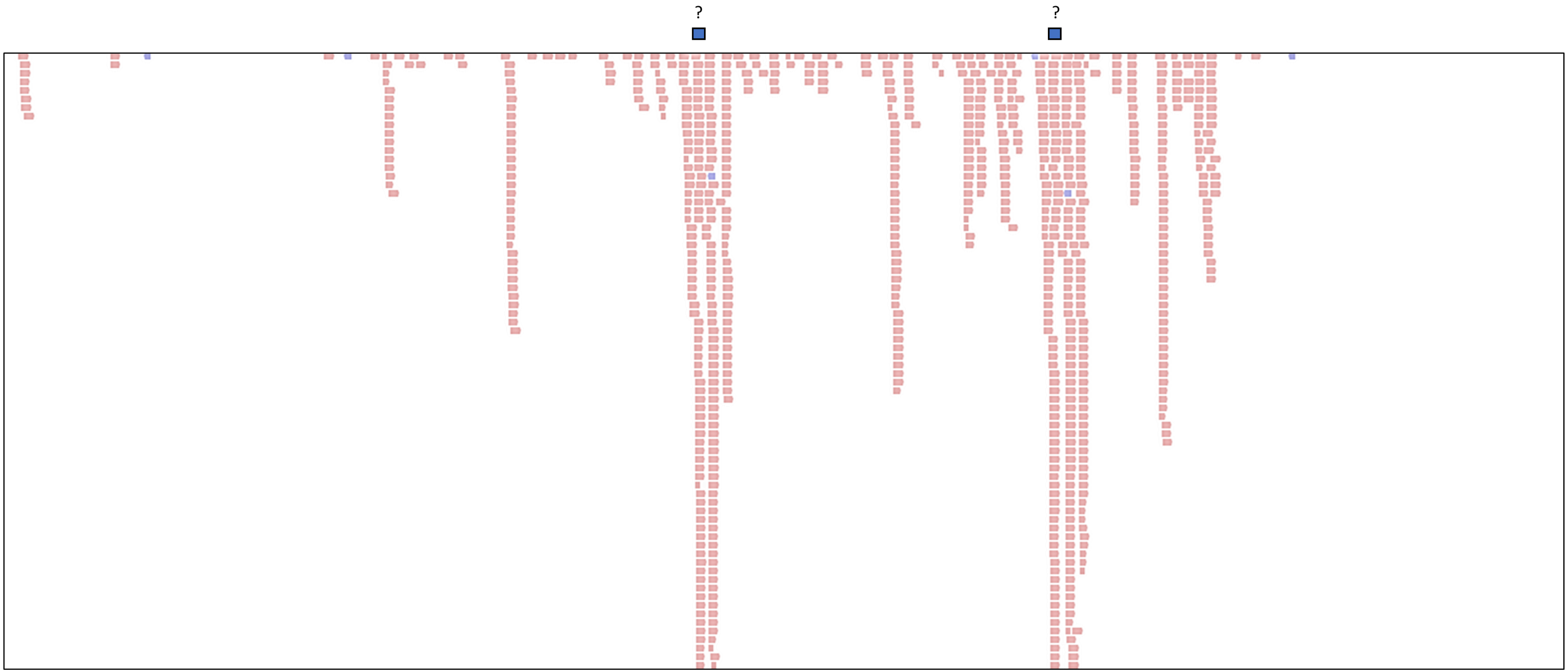


piRNA cluster #3: scaffold367_size99769: 90,134-93,976

### SF4

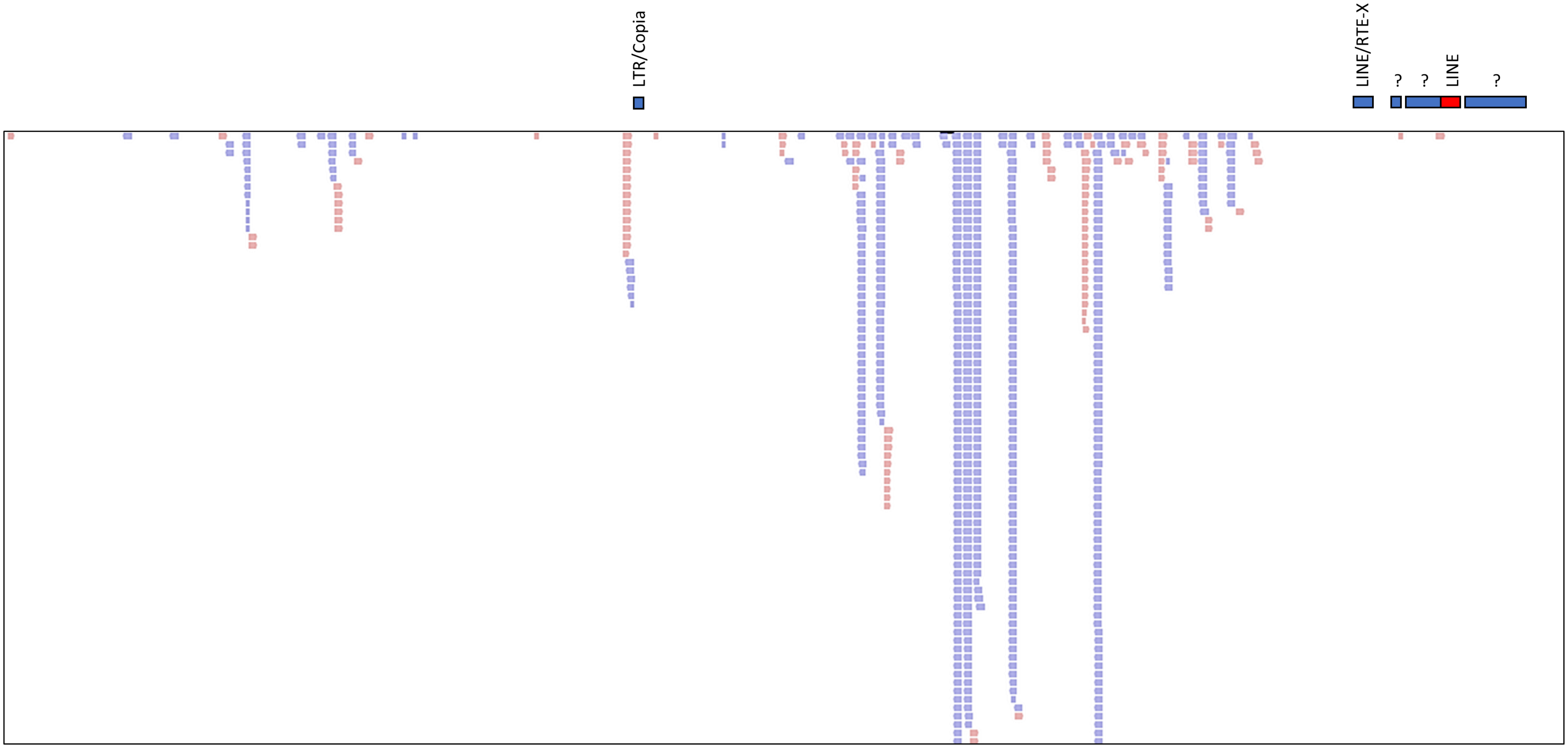


piRNA cluster #4: scaffold386_size93225:37,926-42,144

### SF5

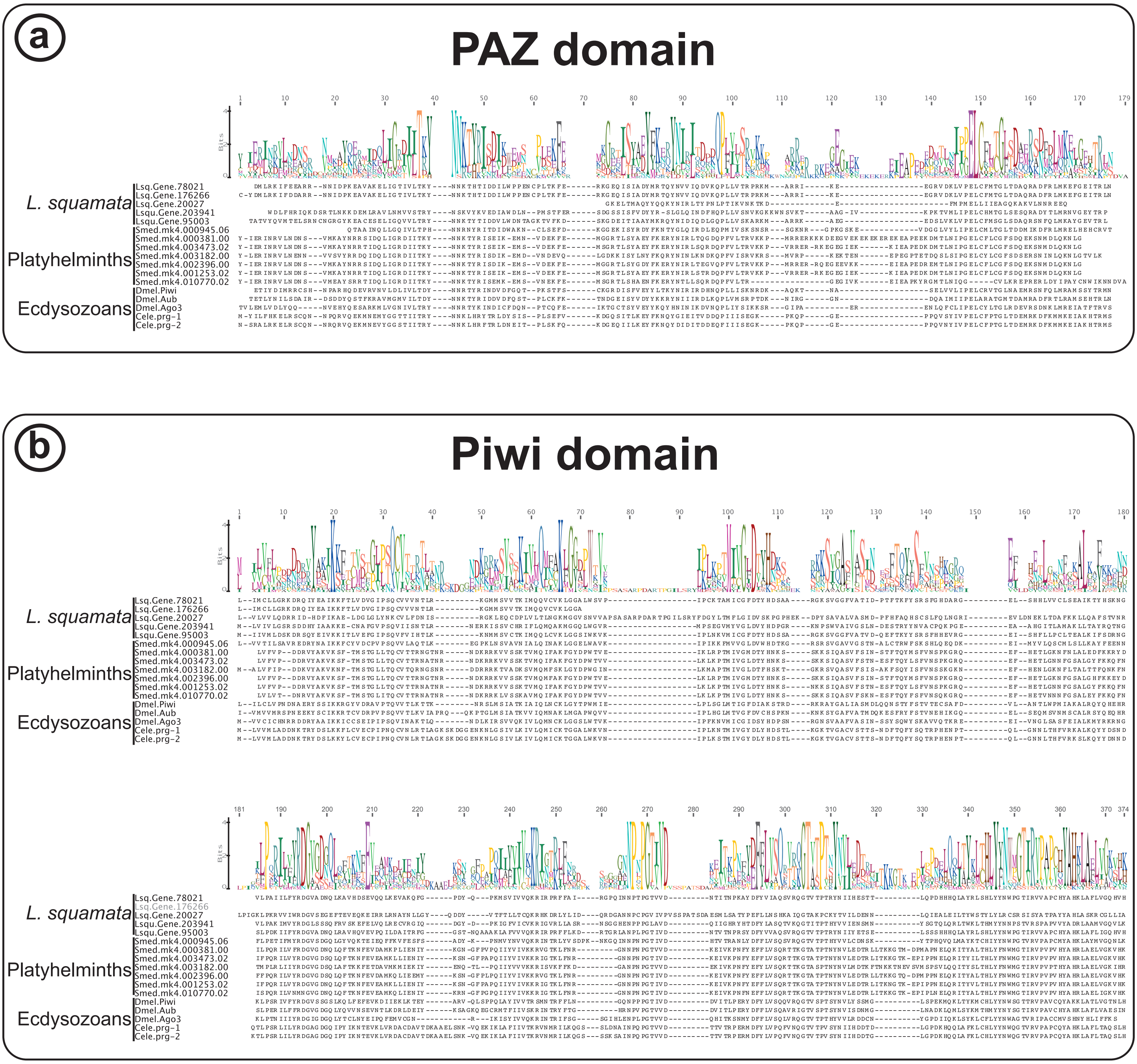
